# Effects of Cannabidiol to circadian period, sleep, life span, close-proximity rhythm, egg reproduction and motor function in *Drosophila melanogaster*

**DOI:** 10.1101/2025.07.08.663811

**Authors:** Haruhisa Kawasaki, Toshihiko Sato, Norio Ishida

## Abstract

Cannabidiol (CBD), a non-psychoactive cannabinoid, has been studied for its various health-promoting effects recently. This study investigates the effects of dietary CBD to the circadian clock of *Drosophila melanogaster* as a model animal and its many physiological effect to flies. We showed that CBD extended the period of locomotor activity in a dose-dependent manner, suggesting its influence on the circadian clock. Additionally, CBD improved sleep quality and extended lifespan under starvation conditions. The study also revealed enhanced rhythmicity in Close Proximity (CP) rhythm and increased eggs reproduction with dietary CBD supplementation. Furthermor, CBD attenuates age-related motor dysfunction in wild-type and Parkinson’s disease (PD) model in *Drosophila*. These findings strongly suggest that appropriate amount of CBD affects the circadian rhythms, sleep, life span, CP rhythm, egg reproduction and motor function of *Drosophila melanogaster*, and providing a basic data for exploring its potential applications in managing circadian-related disorders in other animals.

## Introduction

Cannabidiol (CBD) is a compound derived from the cannabis plant (*Cannabis sativa L.*) and belongs to a group of substances known as cannabinoids. Unlike the primary psychoactive compound in cannabis, tetrahydrocannabinol (THC), CBD is non-psychoactive, non-addictive, and considered a legal substance in many countries. CBD has been reported to induce relaxation and alleviate anxiety and stress (Hazekamp, 2018). Furthermore, it exhibits analgesic and anti-inflammatory properties, making it a promising candidate for managing chronic pain and conditions such as arthritis (Burstein, 2015). CBD is also utilized as a treatment for certain types of epilepsy, including Dravet syndrome and Lennox-Gastaut syndrome (Samanta, 2024) (Devinsky et al., 2017). Its antioxidant properties have also been documented, suggesting potential benefits in reducing cellular damage caused by free radicals and contributing to anti-aging effects (Ambrose & Simmons, 2019). Additionally, it has been associated with improvements in sleep quality, aiding in the management of insomnia and promoting natural sleep cycles (Shannon, Lewis, Lee, & Hughes, 2019).

The circadian clock, often referred to as the biological clock, is an intrinsic timekeeping system that governs a wide range of physiological and behavioral processes in living organisms (Pando & Sassone-Corsi, 2001; Reppert & Weaver, 2001) . This system operates on an approximately 24-hour cycle, aligning internal rhythms with external environmental cues such as the light-dark cycle. In mammals, the central pacemaker of the circadian system is located in the suprachiasmatic nucleus (SCN) of the hypothalamus, which receives photic input from the retina to synchronize bodily functions with the solar day. The circadian clock regulates numerous processes, including hormonal secretion, body temperature, metabolism, close-proximity rhythm and sleep-wake cycles, ensuring optimal physiological performance. For instance, melatonin, a hormone secreted by the pineal gland, promotes sleep during the night, while cortisol, produced by the adrenal glands, facilitates alertness and energy mobilization during the morning (Zisapel, 2018). Disruptions to this finely tuned system, such as those caused by shift work, jet lag, or modern lifestyles characterized by irregular light exposure, have been associated with adverse health outcomes, including metabolic disorders, cardiovascular disease, abnormal menstruation and mental health conditions. Emerging evidence highlights the importance of maintaining circadian alignment for overall health and well-being (Roenneberg & Merrow, 2016). Strategies such as light therapy, time-restricted feeding, and chronopharmacology have been proposed to mitigate circadian disruptions and enhance synchronization between internal and external rhythms. This paper seeks to explore the basic physiology underlying circadian regulation to CBD, the consequences of circadian misalignment, and the potential therapeutic interventions aimed at restoring rhythmicity in affected individuals in the future.

The fruit fly, *Drosophila melanogaster*, is a widely used model organism due to its short generation time, high fecundity, and cost-effective maintenance. With a fully sequenced genome and approximately 75% of disease-related and basic physiology related human genes conserved, *Drosophila* enables effective modeling of various human disorders including Perkinson’s disease (Inoue et al., 2021; Ito, Kawasaki, Suzuki, Takahara, & Ishida, 2017; Nedachi et al., 2025; Su, 2019; Suzuki et al., 2013; Tello, Williams, Eppler, Steinhilb, & Khanna, 2022; Xiu et al., 2022). Its rich genetic toolkit-including RNAi, CRISPR-Cas9, and numerous mutant lines-facilitates precise gene manipulation (Tello et al., 2022). Moreover, *Drosophila* displays behaviors relevant to neuroscience and development, while posing fewer ethical concerns than vertebrate models, making it a practical and versatile system for biomedical research. Though *Drosophila* has many advantages as a model organism for studying the circadian system to affect CBD, significant physiological differences between fruit flies and humans require consideration when extrapolating experimental findings to human biology. Therefore, we avoid making unsupported generalizations beyond what the data justify.

CBD has been considered to have a potential relationship with the circadian clock due to its sleep-improving effects; however, studies specifically investigating the link between CBD and the circadian clock have only emerged recently and remain obscure (Bifulco, Navarra, Laezza, & Pagano, 2020; Lafaye, Desterke, Marulaz, & Benyamina, 2019). Furthermore, research on the effects of CBD in *Drosophila melanogaster* is also limited. In this study, we administered CBD to *Drosophila* to investigate its effects on the circadian clock, as well as to explore its potential health-promoting benefits on reproductive function. In fact, there have been one review examining the effects of CBD on reproductive function, there is no report in *Drosophila* (Cameron et al., 2025). Thus, we want to investigate reproduction, courtship behavior and motor function as indicators of health. This study aims to clarify the effects of CBD on the circadian clock and related physiological functions such as sleep, life span, close proximity rhythm, egg reproduction and motor function in *Drosophila melanogaster*, thereby assessing its potential as a health-promoting compound.

## Materials and methods

### *Drosophila melanogaster* strains and maintenance

The wild-type strain, Oregon-R was raised under a 12-h light/12-h dark cycle at 25°C. All flies were raised in vials with standard fly media consisting of 8% corn meal, 5% glucose, 5% dry yeast extract, and 0.64% agar. Media was boiled and supplemented with 0.5% propionic acid and 0.5% butyl p-hydroxybenzoate as preservatives. For the starvation assay, flies were kept in vials with 1% agar medium. We used male flies for all experiments except for courtship behavior assay. α-synuclein *G51D* is a famous mutation gene for familial Parkinson disease gene in human.(Monteiro Neto, Lima, & Follmer, 2024; Nuber et al., 2024). The *UAS-SNCA-G51D*, and *elav-GAL4/CyO* strains were obtained from the Bloomington *Drosophila* Stock Center (Bloomington, IN, USA). The *IF/CyO*; *elav-GAL4* strain was kindly provided by Dr. Masami Shimoda from The National Agriculture and Food Research Organization (NARO). Transgenic females carrying *UAS-SNCA-G51D* were crossed with males carrying *IF/CyO;elav-GAL4* to generate the *Drosophila* Parkinson’s disease model expressing human *G51D* mutated α-synuclein in neurons (*UAS-SNCA-G51D;elav-GAL4*).

### Preparation of cannabidiol medium

Using a 75mg CBD-containing capsule, CBD4500 (BodyVoice, Chiyoda-ku, Tokyo), a slit was made with a razor blade, and the CBD mixing palm oil was drawn out using a micropipette. The CBD oil was then dissolved in ethanol to prepare the CBD stock solution (100mg/ml). This solution was added to each medium to achieve the desired CBD concentration. All experiments were initiated by placing *Drosophila* in medium containing CBD except for control vials.

### Locomotor and walking distance assays

We tracked the movements of flies that were individually housed with medium, using infrared sensors and a *Drosophila* activity monitor (DAM) system (Trikinetics Inc, Waltham, MA) placed in an incubator under constant dark at 25°C ± 0.5°C for 5 days after 2 days of 12-h light/12-h dark cycle. Signals from the sensors were summed every minute using a computer. Period length was calculated from all data collected using chi-square periodogram analysis (ClockLab, Actimetrics, Inc.).

Walking distance of *Drosophila* was also evaluated by using small-animal behavior analysis system, Automated Circadian Analysis System (AutoCircaS) (Taise Co. ltd. Chiba, Japan) (Inoue et al., 2022). AutoCircaS can trace the movement of flies from consecutive images and analyze the data. Each fly was placed in a 15 mm diameter petri dish with medium containing CBD. Flies image were captured continuously for 5 days under constant dark.

### Assays of sleep behavior

Caffeine (0.5 mg/mL) was used for making insomnia model in fly. Caffeine and CBD were mixed into 5 % glucose-2 % agar medium simultaneously. Male flies were placed individually into 24-well microplates (353047, Corning, NY, USA) containing the medium under diethyl ether anesthesia. Flies were acclimated for 3 days, and then the sleep duration were measured for 3 days by using AutoCircaS (Inoue et al., 2022). AutoCircaS can record the behavior of flies with an infrared CCD camera, analyze the time-lapse images, and automatically calculate the sleep duration. Sleep in flies was defined as a minimum of 5 min of locomotor inactivity (Huber et al., 2004) (Shaw, Cirelli, Greenspan, & Tononi, 2000)

### Life span measurements under starved condition

We determined *Drosophila* survival time under starved condition (Kawasaki, Okano, Ishiwatari, Kishi, & Ishida, 2023; Kawasaki et al., 2021). Lifespan under starvation in 1% agarose was assessed by placing male flies (typically, 60 individuals each) in vials containing 5 mL of 1% agarose at 25 °C under LD 12:12 cycles. Dead flies were counted every few hours (2-14 hours). Statistical data are expressed as means ± SD. Differences in percent survival was statistically evaluated using log-rank test.

### Close-proximity assays

Close-proximity (CP) assays of *Drosophila melanogaster* courtship behavior were performed according to our previous paper (Sakata et al., 2015). At 4-day-old, one male and one female fly of Oregon R strain were anesthetized with diethyl ether and placed in 35-mm-diameter dishes containing the standard or CBD mixed medium. CP index of each couple was measured by using AutoCircaS. The CP index was calculated from the X-Y value with a threshold (<5 mm) between them. Male flies moving to within 5 mm of a female were scored as 1 and those remaining >5 mm from a female were scored as 0 for the algorithm of the CP index program. The circadian rhythmicity of CP was determined using autocorrelation (CORREL function) analysis (Levine, Funes, Dowse, & Hall, 2002). The free-running period and the power of rhythmicity in each genotype were calculated as the average of the free-running period and the maximum correlation between each pair evaluated by autocorrelation as being rhythmic (CORREL function; (Hamasaka, Suzuki, Hanai, & Ishida, 2010). Data were obtained under Light: Dark (LD) 12:12 conditions.

### Measurement of the effective eggs number

Four male and four female flies, two days after emergence, were placed in a vial containing either standard or CBD-mixed medium. Four vials were prepared for each group. They were kept at 25°C for four days to lay eggs, and pupae were counted eight days later.

### Climbing assay

The climbing assay was conducted as described previously (Feany & Bender, 2000; Kawasaki et al., 2023; Kawasaki et al., 2017; Mohite et al., 2018). Groups of 10 male flies per vial (2.5-cm diameter) were gently tapped to the bottom and the number of flies climbing until the 8 cm height within a time period of 10 s was counted. The test was repeated 10 times for each set of flies. CBD was mixed into the food and administered continuously.

## Results

### CBD feeding extends the period of locomotor activity in *Drosophila melanogaster*

To evaluate whether CBD affects the circadian clock in *Drosophila melanogaster*, we measured locomotor activity rhythms using the DAM system. As shown in Figure 1a, the average locomotor activity period in control flies was 23 hours. However, in flies fed with CBD, the locomotor activity period extended in a dose-dependent manner with increasing CBD concentrations (0, 1, 2 mg/mL). Notably, flies that ingested CBD at a concentration of 2 mg/ml exhibited a significantly longer locomotor activity period compared to the control group (p<0.004). The data suggests that CBD feeding extends the period of locomotor activity in *Drosophila melanogaster*.

**Figure 1.**
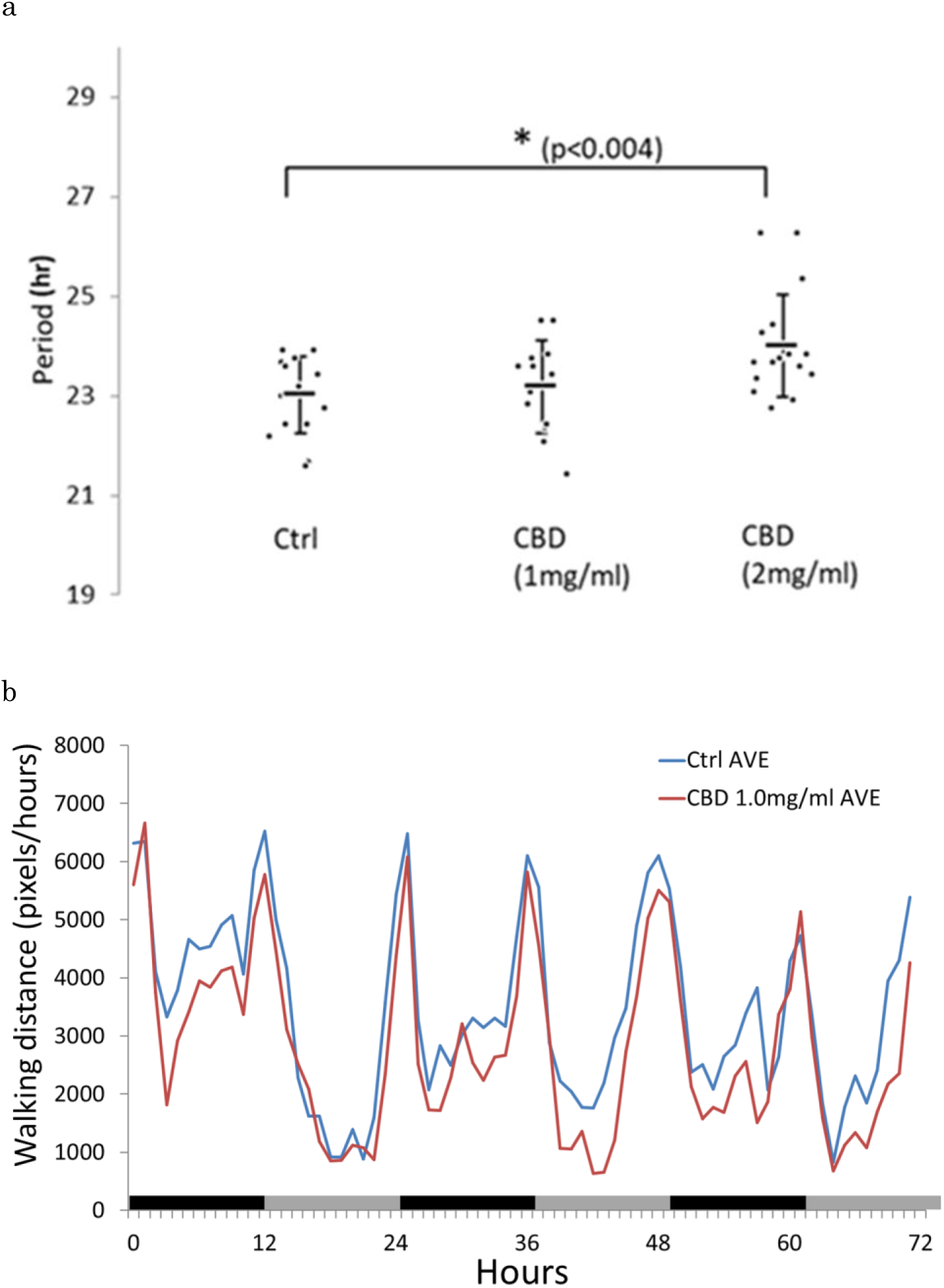
CBD extends the locomotor activity period in *Drosophila melanogaster*. (a) The locomotor activity of wild-type (Oregon R) male flies was assessed using the *Drosophila* Activity Monitoring (DAM) system. CBD was added to the medium at concentrations of 0 (control), 1, and 2 mg/mL. Each dot (circle) represents the locomotor activity period of an individual fly, and squares indicate the mean values. Error bars represent the standard deviation. Approximately 20 flies were used in each group. In the control group, the locomotor activity period was approximately 23 hours. In the 2 mg/mL CBD group, the period was significantly extended. Period length was calculated from all data collected using chi-square periodogram analysis (ClockLab, Actimetrics, Inc.) In the locomotor assay, flies were placed in LD condition for 3day followed by constant dark for 5days. (b) Walking activity of flies (Oregon R; n=18 each) was evaluated using an Automated Circadian Analysis System (AutoCircaS) and represented as waveform data. Each fly was placed in a 15 mm diameter arena with medium containing 0 or 1 mg/mL CBD. The behaviors of flies were time-lapse (one frame per 10 sec) imaged. AutoCircaS recognizes the target as a fly based on pre-set criteria such as size and color intensity. The walking distance of fly is plotted. Data are expressed as the mean. The Y-axis of the graph shows the walking distance of *Drosophila* in pixels.

### Dieting CBD increases sleep in *Drosophila*

As CBD affects the locomotor activity rhythm of *Drosophila melanogaster*, we next conducted sleep measurements using Automated Circadian Analysis System, AutoCircaS (Inoue et al., 2022). The addition of caffeine to the medium (0.5mg/mL) reduced sleep in *Drosophila*, suggesting that insomnia model was established as observed in previous reports (Choi, Ko, Kim, Hong, & Suh, 2017; Inoue et al., 2022; Jo, Jeon, Ahn, Han, & Suh, 2017).

When CBD was further added (0.1 or 0.3 mg/mL) in insomnia model fly, the sleep amount increased in CBD concentration-dependent manner. However, excessive CBD supplementation (1.0 and 3.0mg/mL) led to continuous sleep throughout the day and night (Figure 2a, b).

**Figure 2.**
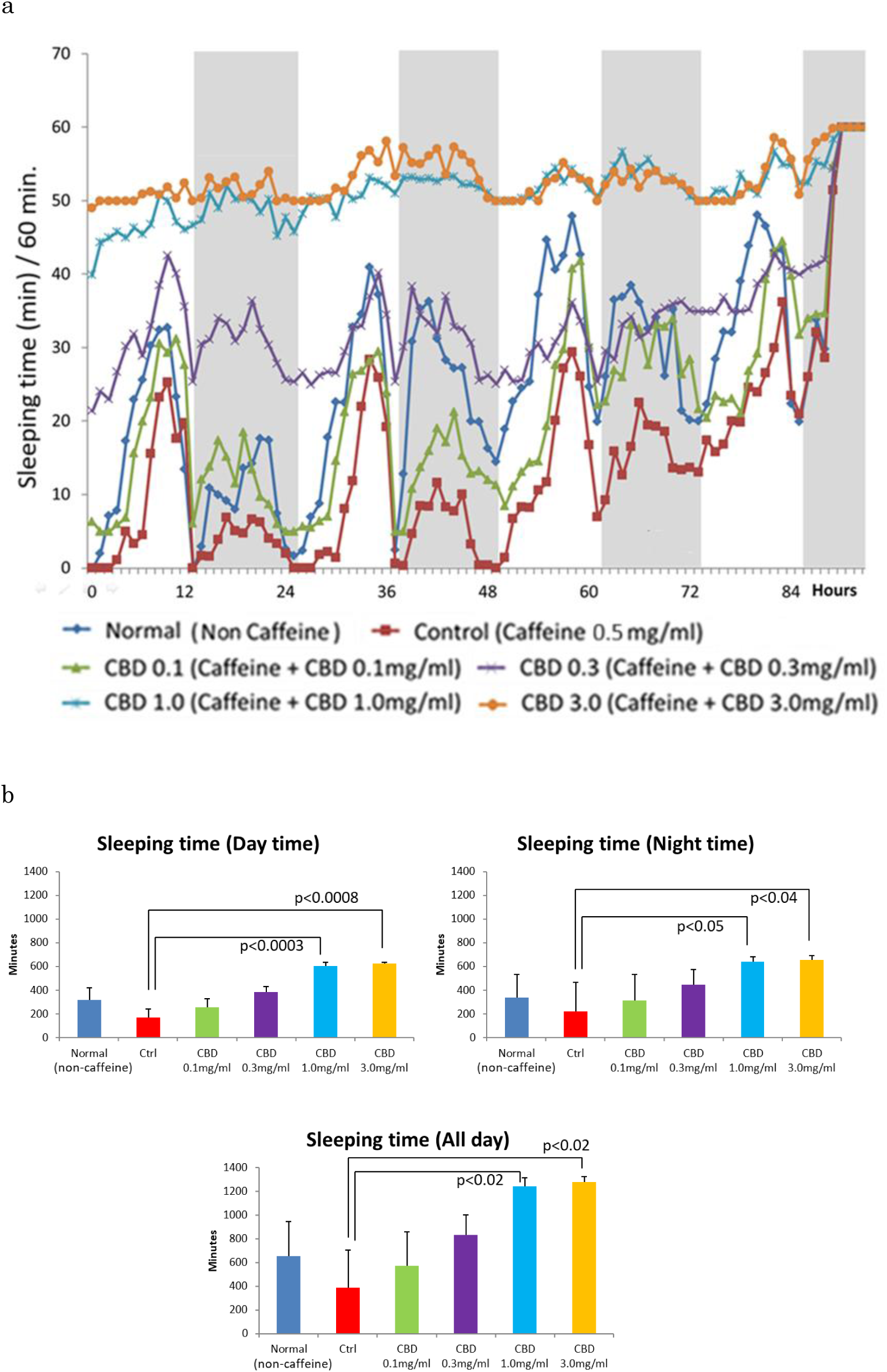
CBD increase caffeine-induced reduction in sleep duration in *Drosophila melanogaster*. (a) The sleep duration of wild-type flies was measured using the AutoSircaS small-animal activity monitoring system and is presented as a line graph (Oregon R; n=12 each). Compared to the normal group (blue line), flies exposed to caffeine exhibited a significant reduction in sleep duration (red line). When CBD was added at concentrations of 0.1 mg/mL (green line) and 0.3 mg/mL (purple line), sleep duration gradually recovered in a dose-dependent manner, approaching normal levels. However, excessive concentrations of CBD, such as 1.0 mg/mL (blue-green line) and 3.0 mg/mL (orange line), resulted in excessively prolonged sleep duration. White/gray areas show the light and dark periods, respectively. (b) Sleep analysis data was presented as a bar graph. The vertical axis shows total sleep. Error bars indicate standard deviation. Asterisk represents significant differences (p < .05, Dunnett test).

### CBD Enhances Survival Under Starvation Conditions in *Drosophila*

To evaluate its potential survival effect, we measured life span of *Drosophila melanogaster* under starvation conditions with CBD (Kawasaki et al., 2021; Liu, Chang, Wu, Lin, & Tsai, 2012). When adult flies were maintained on a non-nutritive medium composed of water and agar to sustain humidity, their population exhibited an S-shaped survival curve, declining steadily and reaching complete mortality within approximately three days. Under these conditions, supplementation of CBD to the medium extended the survival period in a concentration-dependent manner (Figure 3). The data suggest that CBD extends life span of *Drosophila* under starved condition.

**Figure 3.**
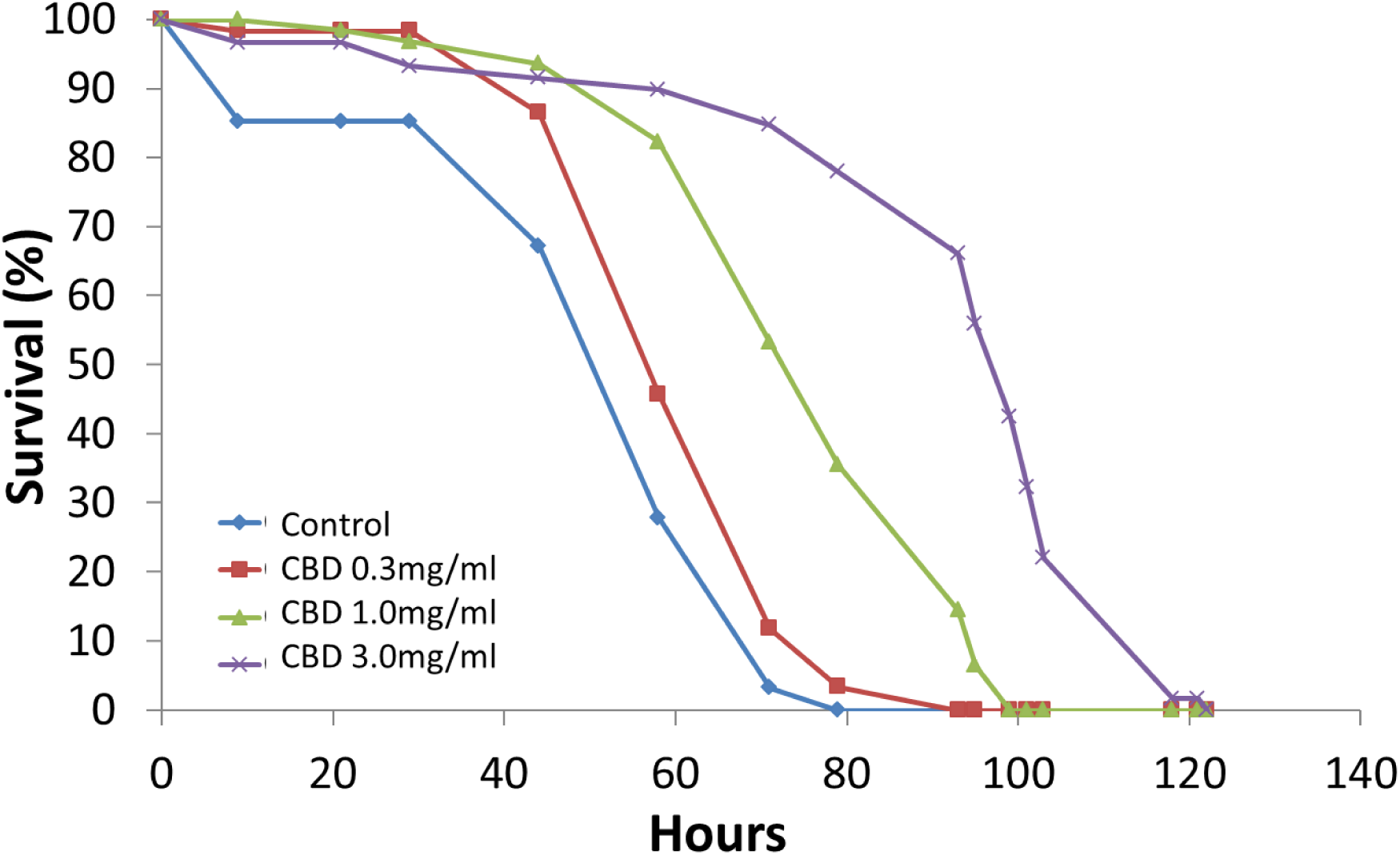
Effects of CBD on the lifespan of wild-type *Drosophila melanogaster* under starvation conditions. Approximately 60 wild-type flies (Oregon R) were used in each group. In the control group (blue line), the number of surviving flies decreased in an S-shaped curve. When CBD was added to the medium at concentrations of 0.3 mg/mL or 1.0 mg/mL, the decline in survival rate was slower (red and green lines, respectively). The results of the log-rank test were as follows: Control vs CBD 0.3mg/ml, p<0.04; Control vs CBD 1.0mg/ml, p<1.75×10^-8^; Ctrl vs CBD 3.0 p<6.34×10^-13^

### CBD increased the amplitude of CP rhythms during day and night in *Drosophila*

As the next indicator of CBD effects, we measured Close Proximity (CP) rhythm. The circadian rhythm of mating succession is controlled by the clock genes, *per* and *tim* in *Drosophila* (Sakai & Ishida, 2001). Heterosexual fly couples exhibit significantly different circadian activity from individual flies, having a brief rest phase around dusk followed by activity throughout the night and early morning (Fujii, Krishnan, Hardin, & Amrein, 2007); The assay is referred to as the close-proximity (CP) rhythm. CP rhythm in *Drosophila melanogaster* exhibits a distinct circadian rhythm and can be analyzed using Automated Circadian Analysis System, AutoCircaS(Inoue et al., 2022). In flies with mutations in clock genes, this rhythm is abolished, indicating that CP rhythm is under clock gene control (Sakata et al., 2015). When we measured the CP rhythm of wild-type flies with fed CBD (0.3 mg/mL and 1.0 mg/mL), the amplitude of CP rhythm increased ie; they were significantly more active during the night time than the day time (Fig 4). This suggests that CBD increased the amplitude of CP rhythms during day and night in *Drosophila.* In contrast, excessive amounts of CBD (3.0mg/ml) had adverse effects in this assay probably because the night time sleep activity unnaturally increased at this concentration(Fig 2a).

**Figure 4.**
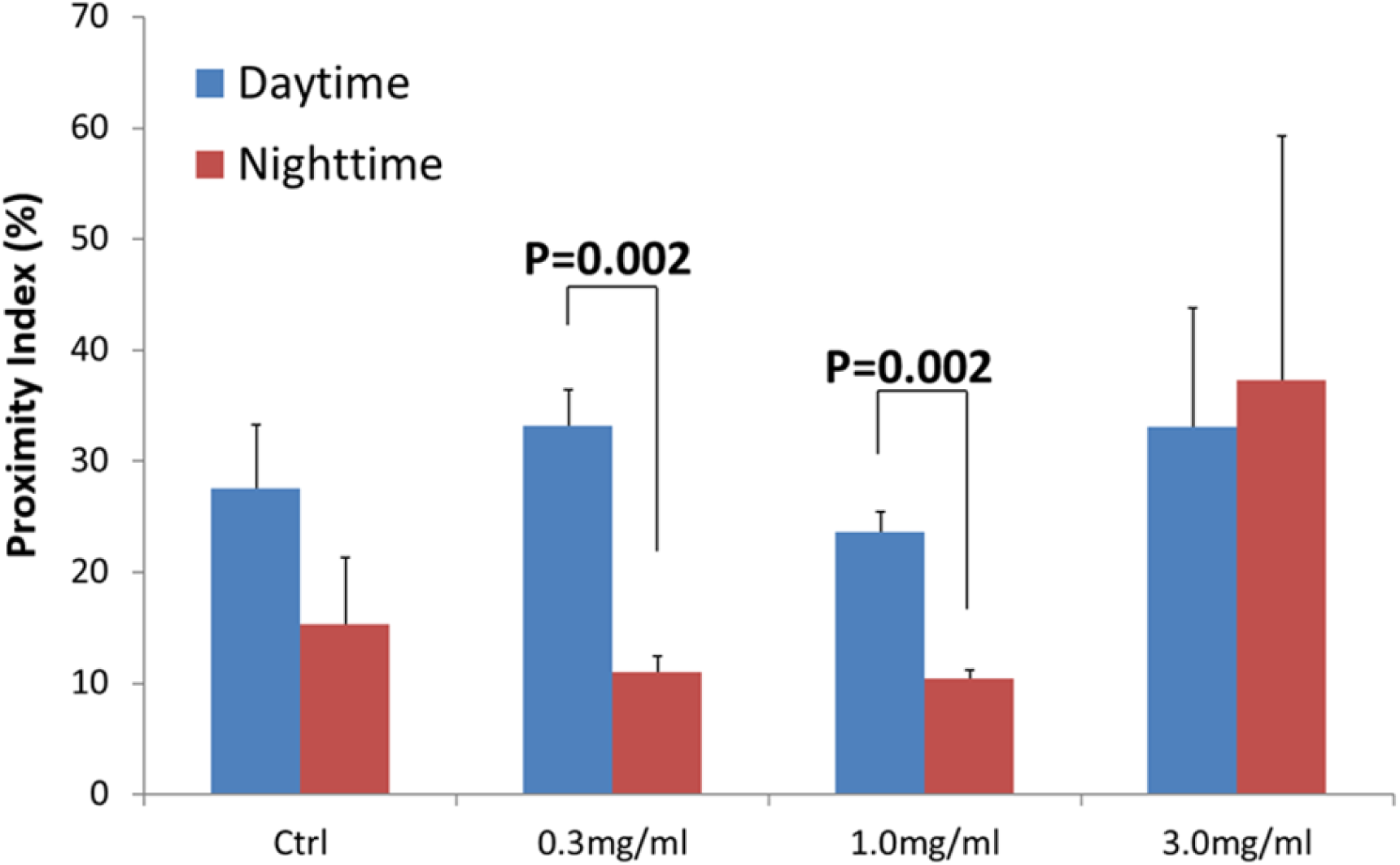
Evaluation of CBD effects on CP rhythm in wild-type *Drosophila melanogaster*. Close-proximity (CP) assays were conducted to evaluate courtship behavior in wild-type flies (Oregon R; 6 pairs each) using the AutoCircaS (Inoue et al., 2022). CP values were analyzed separately for daytime and nighttime and presented as a bar graph. In the control group, CP values were generally higher during the day compared to nighttime. When CBD was added at concentrations of 0.3 mg/mL and 1.0 mg/mL, this diurnal trend became more pronounced, showing significant differences between day and night. However, at excessive concentrations of CBD, this distinction between day and night was diminished. A two-way repeated measures ANOVA showed a significant interaction between CBD administration and time of day (p = 0.007).

### CBD Enhances Successive Egg Reproduction in *Drosophila*

It was observed that CBD affects the courtship behavior of *Drosophila melanogaster.* Next, we evaluated whether CBD influences egg reproduction in flies. To measure the number of viable fertilized eggs laid by female flies, adult flies were removed from vials after four days of egg-laying. After the eggs were developed a few days later, the number of pupae was counted. We refer to this method as “the effective eggs number.” As shown in Figure 5, adding CBD to the medium (concentration of 0.1 mg/mL) increased the effective eggs number compared to the control. The data suggest that CBD Enhances Egg Reproductive Success in *Drosophila*. However, excessive amounts of CBD (1.0 mg/ml) had adverse effects.

**Figure 5.**
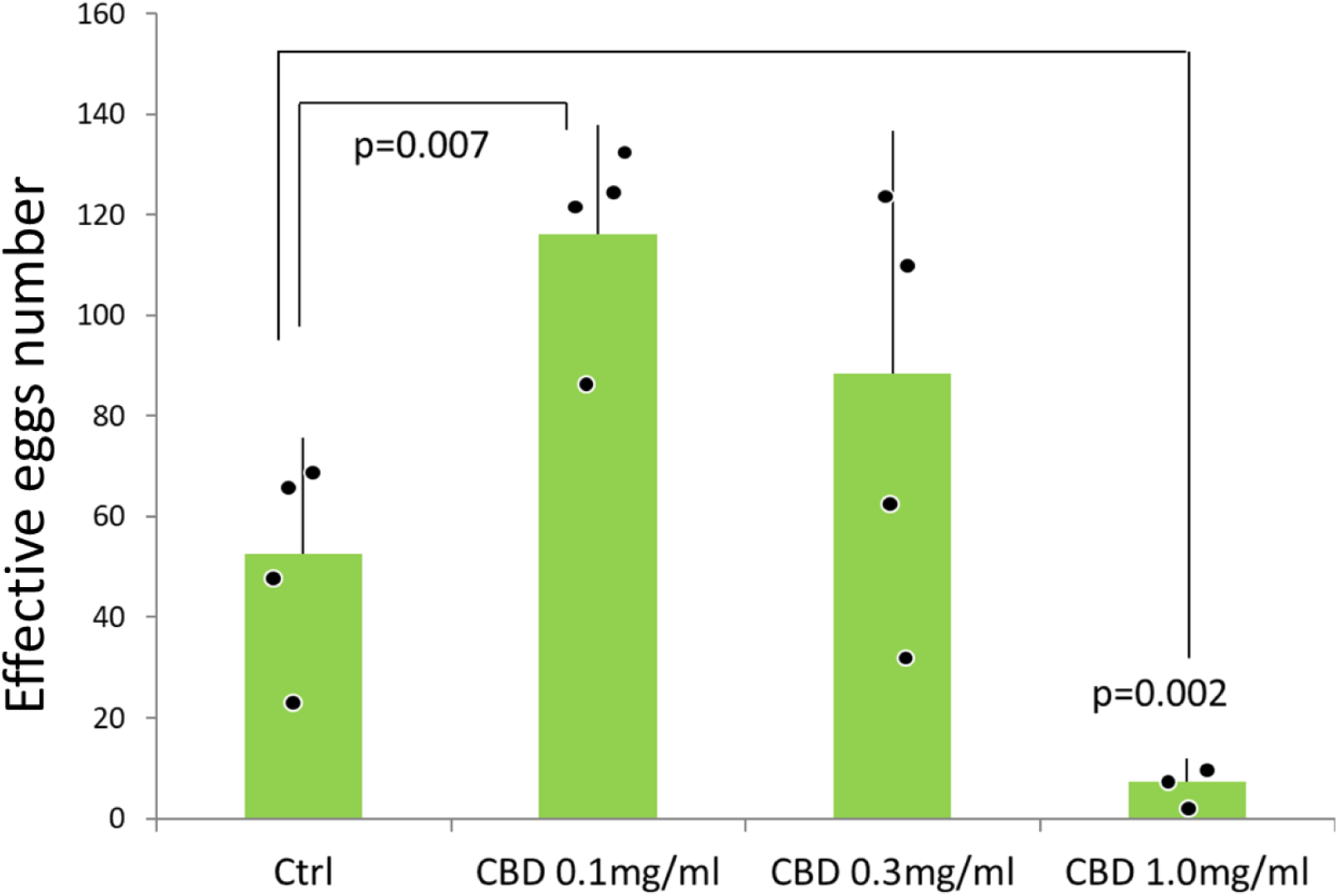
CBD increases the effective eggs number in *Drosophila melanogaster*. Four male and four female adult flies (Oregon R) were placed in a vial containing standard or CBD-mixed medium and allowed to mate freely for 4 days. After removing the adult flies, the vials were maintained at 25°C, and the number of pupae emerging after 8 days was counted as the effective number of eggs laid. Four vials were prepared for each group. In the group where CBD was added to the medium at a concentration of 0.1 mg/mL, the effective eggs number was significantly increased, suggesting that an appropriate amount of CBD promotes egg-laying activity.

### CBD attenuates age-related motor dysfunction in wild-type and Human Parkinson’s disease model in *Drosophila*

Motor function in wild-type and Perkinson’s desease model in *Drosophila melanogaster* were assessed using the climbing assay. Although motor function typically declines with age, administration of CBD containning food reduced this decline in wild-type flies(Figure 6). Remarkably, a similar protective effect of CBD was also observed in a Human Parkinson’s disease model of *Drosophila*, *G51D; elav-gal4*. (Figure 6). The data suggest that CBD attenuates age-related motor dysfunction in wild-type and Parkinson’s disease model in *Drosophila*.

**Figure 6.**
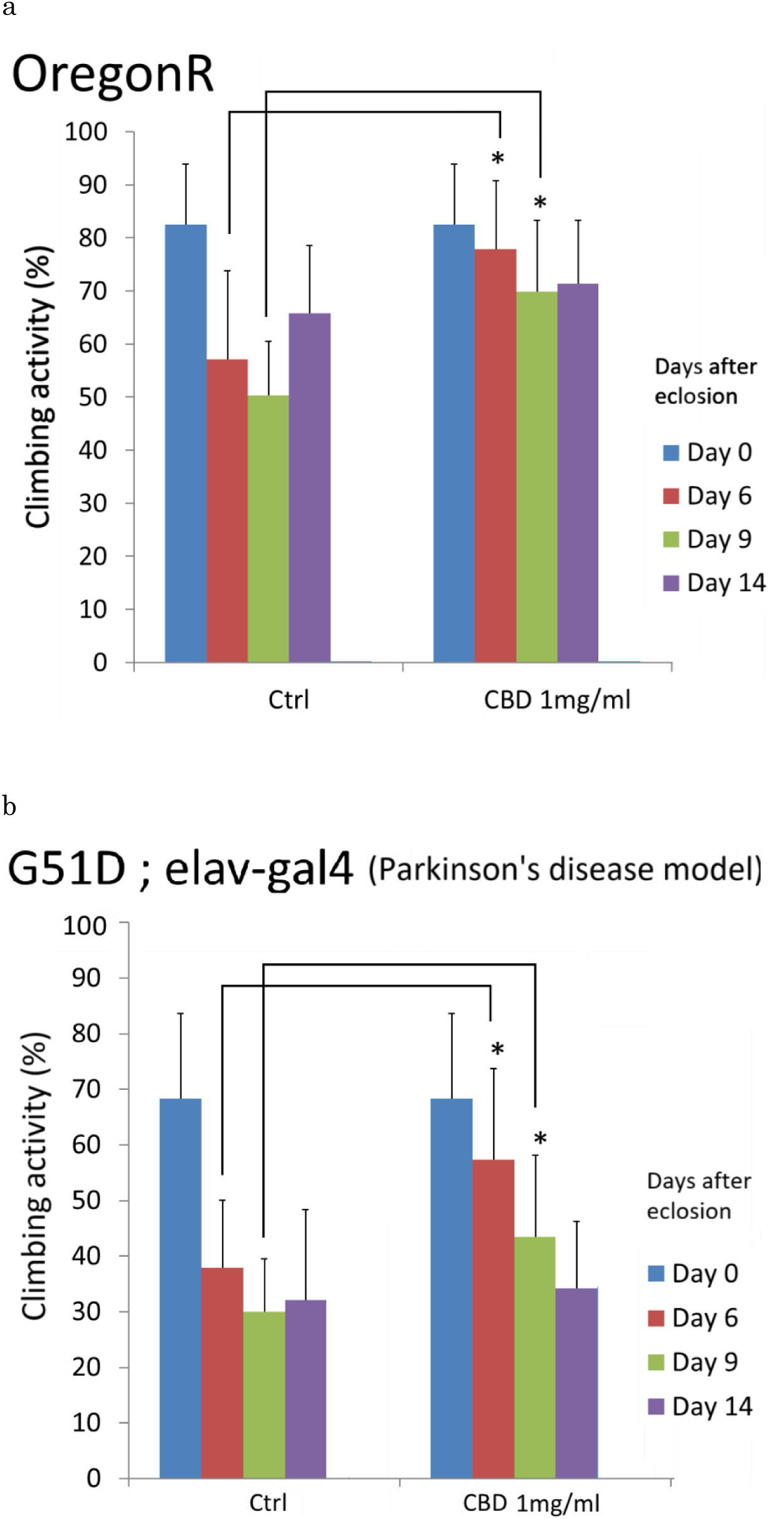
CBD suppresses age-related decline of motor function in wild-type and Parkinson’s disease model *Drosophila*. (a) Climbing activity for wild type (Oregon R) and (b) neurodegenerative disease model flies. G51D is alpha-synuclein *Drosophila* model of Parkinson’s disease. α-Synuclein *G51D* is a famous mutation gene for familial Parkinson gene in human. We compared climbing activity for control and G51D flies. Error bars indicate standard deviation. Statistical data by Student t-test are expressed as mean ± SD. Asterisk represents significant differences (p < .05, t-test).

## Discussion

This study reports novel data that dietary administration of cannabidiol (CBD) alters the circadian rhythm in *Drosophila melanogaster*, resulting in positive health outcomes Including sleep, life span, courtship behavior, egg reproduction and motor function in flies. Research on the effects of CBD in *Drosophila* is extremely limited, with only a few exceptions (Candib et al., 2024; He, Tan, Ng, Rui, & Yu, 2021; Hussain et al., 2024). While numerous studies have explored the relationship between CBD and sleep, to the best of our knowledge, none have directly demonstrated CBD impact on circadian rhythms. There are several reports indicating that injection of agonists targeting CB1, Cannabinoid Receptor 1, an endogenous cannabinoid receptor located in the central nervous system, inhibit light-induced phase delay in mammals (Acuna-Goycolea, Obrietan, & van den Pol, 2010; Niepokny & Mintz, 2023; Sanford, Castillo, & Gannon, 2008). In contrast, a report indicating that CBD injection does not inhibit light-induced phase delays in the circadian rhythms of mammals, even though the administration of CP55,940, a CB1 agonist, suppresses light-induced phase shift in mice (Niepokny & Mintz, 2023). Firstly, our data suggests that CBD affects the period length of the circadian clock. Moreover, our study demonstrates the effects of CBD when administered to animals through oral ingestion rather than injection. To date, precious molecular study has reported that temporal CBD supplementation affected the expression levels of clock genes in cultured mammalian microglial cells (Lafaye et al., 2019); however, the study did not show the data of period of circadian rhythm because lack of continuous monitoring of gene expression. Therefore, our study is the first paper that CBD can lengthen the circadian period in vivo. Thus, the present data suggest that dietary CBD lengthened the period of circadian rhythm in locomotor behavior by using DAM system (Fig1a). We also confirmed the same data using different Automated Circadian Analysis System, AutoCirCas system (Inoue et al., 2022), and interestingly, the addition of CBD to the medium was found to lengthen the period of circadian rhythm in locomotor behavior (Fig1b). In another studies using *Drosophila*, CBD ingestion was reported to extend normal lifespan, but no significant differences were detected in locomotor activity periods, sleep, or starvation experiments compared to the controls(Candib et al., 2024; He et al., 2021).

Our findings indicate that CBD consumption lengthened the locomotor period of *Drosophila* individuals, suggesting its influence on the circadian clock. Lengthening of the circadian period may imply a state of relaxation for human, which could lead to an increase in sleep amount. We firstly showed that oral CBD affects the sleep duration in *Drosophila*. CBD has also been reported to extend the lifespan in zebrafish and *C. elegans* (Frandsen & Narayanasamy, 2022; Pandelides et al., 2020). Thus, enhanced sleep quality and the period of circadian rhythm behavior may contribute to the extension of lifespan in these different species.

Although the precise molecular mechanisms remain unclear, here we showed that CBD administration improved the amplitude of Close Proximity (CP) rhythm in *Drosophila*. In a human context, this may be interesting as promoting the amplitude of rhythmic behavior. In humans, CBD has been reported as a promising therapeutic agent for alleviating the symptoms of endometriosis (Whitaker, Page, Morgan, Horne, & Saunders, 2024). The data suggest that increase of amplitude in rhythmic courtship behavior may be related to this phenomenon. Although we have not yet measured how CBD affects the behavior of fly females, it would be worthwhile to investigate sex difference to CBD in future studies.

Enhanced amplitude of CP rhythm during day time may improve egg reproductive success in *Drosophila*. We showed that CBD intake was associated with a significant increase in the effective eggs number (Fig5). This is very interesting because CBD effect is different between female and male flies. These findings suggest the appropriate amount of CBD may contribute to increasing fertility rates even in mammals. It is known that restricting eating times, rather than eating all day, increases the amplitude of clock gene expression, suppresses obesity, and leads to a healthier lifestyle (Hatori et al., 2012). Similarly, in reproductive activity, clearly distinguishing between day and night and giving reproductive organs sufficient rest may have a positive effect on reproduction. Indeed, several studies have demonstrated that disturbances in the circadian rhythm and sleep negatively affect reproductive function due to endocrine system disorders or oxidative stress (Bahougne, Kretz, Angelopoulou, Jeandidier, & Simonneaux, 2020; Cavalhas-Almeida et al., 2025; Li, Huang, Xu, & Wang, 2024; Miller & Takahashi, 2013; Moralia, Quignon, Simonneaux, & Simonneaux, 2022; Murillo-Rodriguez, Millan-Aldaco, Palomero-Rivero, Mechoulam, & Drucker-Colin, 2006; Yaw, McLane-Svoboda, & Hoffmann, 2020; Yi et al., 2024).

Furthermore, CBD promotes dopamine release, which may underlie its observed effect in suppressing decline of motor function in a *Drosophila* model of Parkinson’s disease. These findings suggest that CBD has the potential to alleviate symptoms of neurodegenerative disorders.

While CBD is well-documented for its health-promoting properties in humans, this study provides evidence that it is one of beneficial compounds for health and is effective material in *Drosophila*, offering insights into its broader applicability for health improvement. However, careful discussion is needed, taking into account interspecies differences between *Drosophila* and mammals. While some studies, such as those by Linares and Schnapp (Linares et al., 2018; Schnapp et al., 2022), reported no significant effects of CBD on sleep in humans, other studies have shown positive effects of CBD on sleep regulation (Chagas et al., 2013; Shannon et al., 2019). The discrepancy between these findings and our study, which observed marked improvements in sleep and behavior in *Drosophila*, may occur from differences in experimental models and conditions. *Drosophila’*s simpler neurological system may cause a more pronounced response to cannabinoids compared to humans. Additionally, variability in human studies—such as differences in participant health, dosage, and experimental design—could explain the contrasting results. These differences suggest that while our findings provide important insights in *Drosophila*, further research is needed to clarify the potential therapeutic effects of CBD on human physiology.

The data strongly suggest that appropriate amount of CBD affects the circadian rhythms, sleep, life span, CP rhythm, egg reproduction and motor function of *Drosophila melanogaster*, and providing a basic data for exploring its potential applications in managing circadian-related disorders in other animals.

## Conclusion

This study provides the first comprehensive investigation of CBD’s effects on the circadian rhythm and basic related physiology such as sleep, lifespan, CP rhythm, egg production, motor function of *Drosophila melanogaster.* The extension of the period of locomotor activity and improving sleep quality suggest that CBD can lengthened the circadian clock. Furthermore, the increased survival under starvation, enhanced courtship rhythmicity, improving the reproductive success and improved motor function in PD model highlight the broad-spectrum health-promoting effects of CBD. The results provide a basic data for future research on the applications of CBD in controlling circadian disruptions and related health issues, potentially extending to human aging and veterinary medicine.

